# Bidirectional chemogenetic modulation of claustral activity causes altered cortical dynamics

**DOI:** 10.1101/2023.11.07.566057

**Authors:** Ryan Zahacy, Yonglie Ma, Ian R. Winship, Jesse Jackson, Allen W. Chan

**Affiliations:** Neuroscience and Mental Health Institute, University of Alberta, Edmonton, Alberta, Canada; Department of Psychiatry, University of Alberta, Edmonton, Alberta, Canada; Department of Physiology, University of Alberta, Edmonton, Alberta, Canada

**Author notes:** Co-senior author and correspondence, or.

**Keywords:** claustrum, calcium imaging, mouse, cortex, functional connectivity, spontaneous activity, resting-states, mesoscale, chemogenetics

## Abstract

In recent years, there has been a growing interest in understanding the function of the claustrum. The claustrum is a thin, long, subcortical structure with dense connections to the cortex. Despite these extensive connections, the manner in which the claustrum influences broader cortical activity remains unclear. We used mesoscale calcium fluorescence imaging to examine resting state cortical activity (1-5Hz) and sensory-evoked responses in lightly anesthetized mice, before and after bi-directional chemogenetic modulation of claustrum-prefrontal neurons. Claustrum inhibition resulted in increased activity in anterior-medial cortical areas, whereas claustrum excitation resulted in decreased activity in anterior-medial cortical regions. Claustrum inhibition also led to an increased local coupling between areas of frontal cortex, a decreased correlation between anterior medial regions-of-interest (ROIs) and lateral and posterior ROIs, and an increased sensory evoked response in the visual cortex. These results are consistent with the finding that the claustrum has a large feed-forward inhibitory effect on the PFC, and the concept that specific claustrocortical pathways may modulate recruit and synchronize activity in topographically distinct cortical modules. Together these results show that neural activity in the claustrum modulates the excitability of prefrontal cortical networks, suggesting a potential target for prefrontal-dependent behaviors such as learning, attention, and stress regulation.

## Introduction

The claustrum is a subcortical structure identified in all mammals that have been examined (Wang et al., 2017), as well as in the Australian bearded dragon (*Pogona vitticeps*) and turtle (*Trachemys scripta*) (Norimoto et al., 2020). Its unique features, such as its long thin structure and dense connectivity with the cortex (Torgerson et al., 2011, Wang et al., 2017, Wang et al., 2023), have led to theories proposing a function for the claustrum in selective attention (White et al., 2020; Atlan et al., 2018), impulsivity (Liu et al., 2019), stress (Niu et al., 2022), sensory detection (Smith et al., 2019, Remedios et al., 2010), and decision making (Chevee et al., 2021). Numerous studies have explored the anatomical and physiological organization of the claustrum with different cortical subregions (Wang et al., 2017; Chia et al., 2020; Marriott et al., 2020), and collectively these works have supported integration of the claustrum within several cortical networks, including the default mode network and salience network, based on its dense connectivity with major network hubs (Smith et al., 2019, Qadir et al., 2022). However, an understanding of how claustral neural activity modulates neural activity in the cortex remains unclear.

Previously it has been shown that claustrum activation has a dissociable influence on activity in different cortical regions. Projections to the prefrontal region generate strong feed-forward inhibition, whereas inputs to posterior regions, such as the retrosplenial cortex, are more likely to generate both inhibition and activation of cortical excitatory cells (Jackson et al., 2018; Narikiyo et al., 2020; Peng et al., 2021; McBride et al., 2023; de la Torre-Martinez et al., 2023). Claustrum outputs are known to exhibit two main juxtaposing features. While some claustrum neurons exhibit a broad pattern of axonal collateralization across several cortical regions (Wang et al., 2017; Narikiyo et al., 2020; Wang et al., 2023), there is a topographically organization wherein the dorsal/ventral claustrum projects to anterior/posterior cortical regions respectively (Smith and Alloway, 2014; Marriott et al., 2020; Wang et al., 2023). Very limited work has been done to assess how subpopulations of claustrum neurons modulate activity in different cortical targets. Given that the claustrum has broad and specific pathways to communicate with the cortex, it is unclear if manipulating a subpopulation of claustrum projections will lead to widescale changes in cortical activity or more localized changes within a dedicated cortical center (Torgerson et al., 2011; Wang et al., 2017; Wang et al., 2023). We previously demonstrated that the claustrum provides strong, widespread, and long-lasting feedforward inhibition of the prefrontal cortex (PFC) sufficient to suppress ongoing neural activity. The prefrontal cortex’s vital role in information processing is well-established, with its extensive connections to various cortical and subcortical regions indicating its centrality as an information network hub (Haber et al., 2021; Le Merre et al., 2021; Gao et al., 2022). As a result, modulation of PFC activity through this pathway may facilitate its function by synchronization or integration of its activity with broader cortical networks. However, the comprehensive modulatory effects of claustrum inputs on the broader PFC function and related cortical landscape remain unclear.

Novel imaging techniques such as mesoscale imaging are highly suited for simultaneously measuring cortical activity and functional connectivity across distant areas with high spatiotemporal resolution (Ferezou et al., 2007; Mohajerani and Chan et al., 2013; Oh et al., 2014; Chan and Mohajerani et al., 2015; Ma et al., 2016; Vanni et al., 2017; Bauer et al., 2018). We performed mesoscale cortical calcium imaging in mice expressing GCaMP6s (Chen et al., 2013), while chemogenetically activating or suppressing claustrum outputs to the prefrontal cortex. The goal was to determine if pathway-specific modulation of claustrum activity results in changes in cortical activity that were restricted to the prefrontal cortex, or if the modulation of activity extends beyond the principle claustrocortical pathway being manipulated. Our results show a selective modulation of prefrontal activity by claustrum manipulation, while also changing the functional connectivity across prefrontal and posterior cortex. The data, therefore demonstrate that the claustrum can modulate the long-range coupling of cortical networks through its connectivity with the prefrontal cortex.

## Materials and Methods

All procedures are conducted in accordance with the Canadian Council for Animal Care and are approved by the University of Alberta Health Sciences Laboratory Animal Services Animal Care and Use Committee.

### Animals

Adult (25-33 g) C57BL/6J-Tg(Thy1-GCaMP6s)GP4.3Dkim/J (JAX stock #024275) mice aged 4-12 months, of both sexes, were used for all experiments (n=18). These mice express the intensity-based, genetically encoded calcium indicator, GCaMP6s, under the *Thy1* promoter, exhibiting widespread and uniform expression throughout cortex (Chen et al., 2013; Dana et al., 2014). Mice were group housed in cages of 3-5 mice in climate-controlled individually ventilated cages, with a 12-hour light/dark cycle. Mice were given *ad libitum* access to water and standard laboratory mouse diet at all times.

### Surgery

Cortical imaging was performed through a surgically implanted, chronic, transcranial window as described previously (Silasi et al., 2016; Vanni et al., 2017; Xiao et al., 2017). Animals were anesthetized in an induction chamber with isoflurane (4%) and kept at maintenance for surgery between 2-2.5% isoflurane with pure oxygen as the carrier gas. Ophthalmic gel was applied to the eyes to maintain moisture during the procedures, and body temperature was monitored and maintained at 37 ^°^C using an electric heating pad with a feedback thermistor. Bupivacaine (0.25%) was injected into the scalp prior to the initial incision. For head-plate implantation, the scalp of the mouse was removed, and the skull was cleared using a scalpel. Tissue was removed to accommodate a hexagonal head-plate which encased an approximately 10 x 10 mm glass coverslip. Glass coverslips and head-plates were secured to the cleaned skull with transparent dental cement (C&B – Metabond, Parkell Inc. Edgewood, NY, USA). Following all surgeries, animals were treated with buprenorphine (0.1 mg/kg) injected subcutaneously and placed in a recovery cage with a Heat Therapy Pump HTP-5000 (Kent Scientific Corporation, Torrington, CT, USA) water heater.

### Viral Injection

Chemogenetic interventions such as Designer Receptors Exclusively Activated by Designer Drugs (DREADDs) allow for selective transient modulation of neural activity via injection of synthetic agonists (Armbruster et al., 2007). Mice were injected with AAV5-DIO-hM4Di-GFP or AAV5-DIO-hM3Dq-mCherry to induce inhibition or excitation in the claustrum, respectively. To perform the intracranial injections, small burr hole craniotomies were created using a high-speed dental drill and Cre-dependant HM4Di/HM3Dq DREADDs were bilaterally into the claustrum, and a retro-Cre was injected bilaterally into the PFC using a hydraulic microinjector. For injections, anterior-posterior (A/P) and medio-lateral (M/L) coordinates were measured relative to the Bregma, and dorsal-ventral (D/V) coordinates were measured from the surface of the brain for each injection location. Injection coordinates were as follows: PFC: A/P +1.7mm, M/L 0.5mm D/ V, −0.7mm; Claustrum: A/P +1.25mm, M/L +/-2.5mm, D/V −2.5mm. Mice were injected with 200 nL of retro-Cre or DREADD at each location injection at a rate of 100 nL/min, with the glass pipette being left in position for 5 minutes following each injection. Clozapine N-oxide (CNO) was injected at 5 mg/kg to activate the DREADDs. As CNO can be reverse metabolized to clozapine *in vivo* in mice and rats (Manvich et al., 2018), we used mice not infected with DREADDs but injected with CNO as control animals to ensure the effects were not due to the CNO injection.

### Data acquisition

#### Wide Field Optical Imaging

Wide-field optical imaging was performed to capture the global cortical changes in calcium fluorescence as an indicator of neural activity. Calcium fluorescence data were collected as 12-bit images captured with a Teledyne DALSA (Waterloo, ON, Canada) 1M60 CCD camera and EPIX EL1 frame grabber with XCAP 3.8 imaging software (EPIX, Inc., Buffalo Grove IL). GCaMP6s was excited with an LED (470 nm, Thor Labs (Newton, New Jersey, USA) M470L2-C2) filtered through a 470 ±10 nm excitation filter (Thor Labs (Newton, New Jersey, USA), FBH470-10). Calcium fluorescence was filtered using a Semrock (New York, NY, USA) Brightline FF01-525/30-25 green light emission filter housed in a 3D printed filter housing (**Figure 1A**).

**Figure 1.**
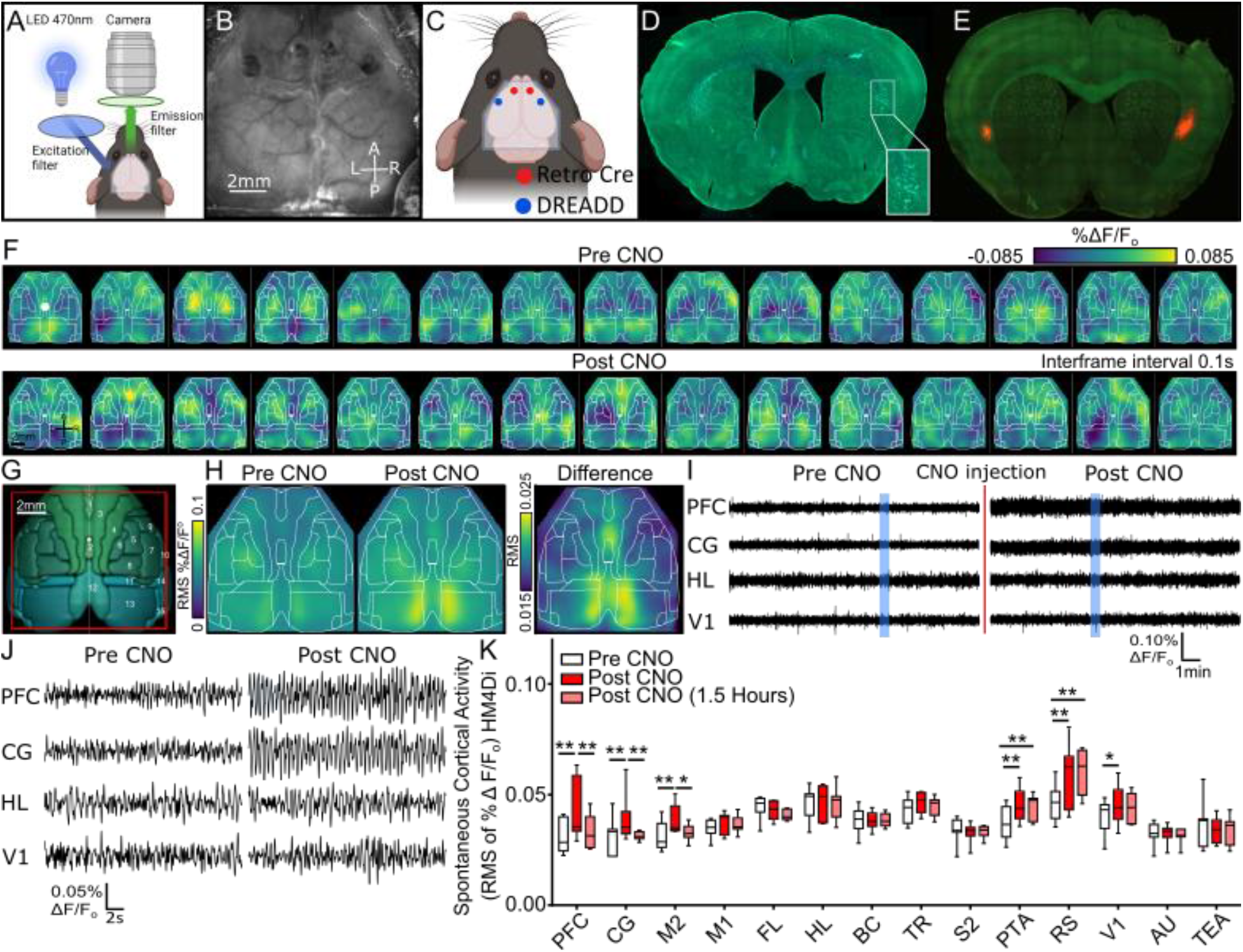
Chemogenetic inhibition of claustrum activity with HM4Di causes increased spontaneous cortical activity. **A)** Schematic diagram of imaging apparatus. Created with BioRender.com **B)** Example of imaging field-of-view revealing the surface of the dorsal neocortex through glass coverslip. **C)** Parcellated region-of-interest (ROI) map showing DREADD (blue) and retro-Cre (red) injection locations. Created with BioRender.com **D)** Coronal brain sections illustrating region-specific expression of DREADD injection locations indicated by GFP (see inset) DAPI (4′,6-diamidino-2-phenylindole) labelling of cell bodies. **E)** Coronal brain section illustrating DREADD injection locations indicated by fluorescent mCherry. **F)** Montage of exemplar spontaneous cortical activity before (Pre) and after (Post) CNO injection (5 mg/kg i.p.). Inter-frame interval of 0.1 s, white dot denotes the bregma. **G)** Schematic of cortical regions of interest based on Allen Brain Institute 3D Brain Explorer, red box approximates imaging field of view. **H)** Mean cortical activity map of calculated root-mean-square (RMS) values across animals for each pixel on in imaging field of view before (left) and following (right) CNO treatment (5 mg/kg i.p.), n=7. Right panel shows the change in spontaneous activity (RMS) between the 2 conditions, where higher values indicated greater increases in RMS **I)** Time course of a 15 minute recording from PFC, CG, HL, and V1 showing stable spontaneous activity over the duration of recording. Shaded blue areas indicate regions expand in Figure 1J. **J)** Example 30 s time-courses of spontaneous activity from PFC, CG, HL, and V1 before (Pre) and following (Post) CNO injection. **K)** Summary of changes in spontaneous cortical activity (n=7). Boxplot of RMS values calculated from the spontaneous cortical activity time course of the calcium signal for 14 ROIs spanning the entire cortical hemisphere. For each ROI, mean activity values were calculated for recordings prior to (white), immediately following (red), and 1.5 hours following (light red) CNO treatment (5 mg/kg i.p.). Statistical significance was calculated using a two-way ANOVA with a Tukey Test to correct for multiple comparisons. * P ≤ 0.05, ** P ≤ 0.01. Two-Way ANOVA with multiple groups comparison.

#### Recordings of spontaneous and evoked activity

Recordings were acquired from mice anesthetized with isoflurane (1-1.25%). For each animal, a baseline spontaneous recording, and sensory evoked recordings were performed (Pre CNO). After these recordings, CNO was administered via intraperitoneal injection (5 mg/kg). Ten minutes following CNO administration, another spontaneous recording and round of sensory stimulation recordings were performed (Post CNO). The final recording of each session was a third spontaneous recording approximately 1.5 hours after CNO administration (Post CNO 1.5 hours).

Calcium fluorescence imaging of spontaneous activity was acquired at 30 Hz in a dark, enclosed chamber and in the absence of visual, olfactory, tactile, or auditory stimulation. As animal brain states exhibit spontaneous change, sensory-evoked recordings were averaged 20 trials of stimulus presentation to reduce these effects. A 10-second interval between each sensory stimulation was used. To reduce potential fluorescence signal distortion caused by the presence of large cortical blood vessels, we focused the camera into the cortex.

For sensory-evoked activity recordings, imaging data were collected at 10 Hz. Sensory-evoked recordings consisted of trials of 20 second recordings with an additional 10 second unrecorded inter-trial interval, and stimulation delivered 3 seconds into each trial. Trials and stimulation were initiated by a signal from a Cambridge Electronic Design 1401 mk2 output board controlled by the software Spike2 (v6.18) (Cambridge, United Kingdom). Stimulation signals were generated using a Model 2100 Isolated Pulse Stimulator (AM Systems, Sequim, WA, USA). Visual stimulation was produced using an LED driver that powered a blue LED light (Rebel LED SinkPAD-II 20 mm Tri-Star, 470nm, Luxeon Star LEDs Quadica Developments Inc., Lethbridge, Alberta, Canada) for a 5 ms directed to the eye. Somatosensory stimulation was generated using a piezoelectric actuator placed in contact with a limb or whisker. Tactile stimulation consisted of a 400 ms stimulus at 10 Hz. Stimulation trials were only included in analysis for animals that had responses for both the Pre CNO and Post CNO conditions.

#### Data Analysis

Images acquired of spontaneous cortical activity were bandpass filtered (1-5 Hz) using a Two-way Chebyshev bandpass filter, a consistent with previous calcium fluorescence mesoscale imaging work (Vanni et al., 2017, Wright et al., 2017). Non-cortex regions of the imaging field-of-view were masked out and recordings were processed with a global signal regression to remove global changes, ΔF/F˳ was calculated to determine relative changes in fluorescence (Mohajerani and Chan et al., 2013; Chan and Mohajerani et al., 2015; Haupt et al., 2017; Vanni et al., 2017). Region-of-interest (ROI) locations were derived from experimentally mapped cortical regions and extrapolations from the Allen Mouse Brain Atlas, and ROIs were represented as 5x5 pixel boxes. For spontaneous activity recordings, frames that exhibited values exceeding the mean ± twice the standard deviation, were removed. As chemogenetic manipulation was bilateral, we anticipated both hemispheres to respond to CNO injections similarly. To confirm this, we compared the root-mean-square (RMS) values from each hemisphere for each condition using a two-way ANOVA with Bonferroni’s Test to correct for multiple comparisons, which showed no ROIs having significant differences (**Supplemental Figure 1**). As such we averaged right and left hemisphere ROI RMS values for each mouse to calculate mean RMS values (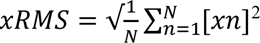; MathWorks Inc., 2022).

#### Perfusion

Animals were anesthetized with isoflurane and subsequently urethane (4%) prior to perfusion. Animals were perfused using first saline, followed by formalin (4%) to fix the brain. The brains were placed in formalin after extraction. Brains were sliced using a Leica Bio Systems (Wetzlar, Germany) VT1000s vibratome. Brains injected with hM4Di were stained with 4′,6-diamidino-2-phenylindole (DAPI) to identify gross cell morphology, and slices were imaged using a Zeiss Axioscan.Z1 slide scanner.

#### Seed pixel Correlation Maps

As previously, seed pixel correlation maps were created by computing the pixelwise correlation coefficients between the time-varying spontaneous activity from the ROI pixel and every other pixel within the image (Mohajerani and Chan et al., 2013; Chan and Mohajerani et al., 2015; Vanni et al., 2017). To quantify the area of local functional connectivity, we calculated area-specific thresholds for the seed pixel of each ROI to set a consistent Pre CNO area, restricted in space to the right hemisphere (**Figure 6B**). We then used that same threshold for the Post CNO recordings for each mouse to calculate relative area changes. Other thresholds were tested that showed similar trends but with lower thresholds the areas were not hemisphere-specific (**Supplement Figure 3**).

### Statistical Analysis

Statistics were done using MATLAB (Mathworks, Natick, MA, USA) and GraphPad Prism software. Analysis of spontaneous activity (RMS) was calculated using a two-way ANOVA with Tukey’s Test to correct for multiple groups comparison. Relative changes in spontaneous activity (RMS) were calculated using (Post CNO -Pre CNO/Pre CNO)*100, and a two-way ANOVA with Dunnett’s Test to correct for multiple groups comparison was used to compare the relative change between the DREADD and control conditions. In box plots whiskers indicate max and min ranges, box 25^th^-75^th^ percentiles, and the line in the box indicates median value. The change in peak sensory response values for Pre and Post CNO conditions were compared using paired t-tests. Error bars and ± represent standard error of the mean. *, and ** indicates P<0.05, P<0.01 respectively.

Seed pixel area calculations were based on a Pre CNO area threshold specific to each ROI. Area of the Post CNO condition was calculated based on the Pre CNO animal-specific threshold for each ROI and calculated as a relative percent change. These values were compared to the standardized baseline using a one-sample t-test. Correlation matrix groups were compared using a Wilcoxon rank sum test in MATLAB (MathWorks Inc., 2022).

### Correlation Matrices

To normalize correlation matrices and facilitate between-individuals comparisons, we transformed Pearson correlation coefficient values to z-scores for each correlation matrix as described previously (Veronese et al., 2019; Yeh et al., 2020). We then averaged the z-scored matrices across animals for Pre CNO and Post CNO conditions. To better visualize functional connectivity changes we clustered related regional functional areas. Primary somatosensory cortices (FL, HL, BC, TR) showed similar patterns of correlation, as such we grouped them into a single somatosensory group (SOM) for correlation matrix analysis.

To quantify the difference between Pre and Post CNO matrices we grouped ROIs into 2 clusters, an anterior-medial region that included PFC, CG, M1, and M2, and a posterior-lateral region which included the other ROIs (S2, PTA, RS, V1, AU, TEA, and SOM). For clarity, we removed self-correlated elements along the diagonal of the correlation matrix and also only display the lower triangle of values as these element values are symmetrical along the diagonal axis (Connor et al., 2016; Vanni et al., 2017). We calculated cumulative distribution functions (CDFs) comparing anterior-medial vs anterior-medial, anterior-medial vs posterior-lateral, posterior-lateral vs posterior-lateral connections between the baseline and Post CNO conditions and evaluated the differences using a Wilcoxon signed-rank test (McGirr et al., 2017; MathWorks Inc., 2022).

## Results

### Spontaneous cortical activity

To measure spontaneous cortical activity, we performed mesoscale, wide-field calcium fluorescent imaging to assess widespread cortical activity with high spatiotemporal resolution (Ferezou et al., 2007; Mohajerani and Chan et al., 2013; Oh et al., 2014; Chan and Mohajerani et al., 2015; Ma et al., 2016; Vanni et al., 2017; Bauer et al., 2018). As previously, targeted manipulation of claustrocortical circuits was attained by the injection of Cre-dependant chemogenetic AAVs bilaterally into the claustrum and AAV-retro-syn-Cre bilaterally into the PFC (Jackson et al., 2018) (**Figure 1D, Methods and Materials**). This allowed for selective inhibition (hM4Di) or excitation (hM3Dq) of claustrum neurons projecting to the PFC. Claustrum specific targeting was confirmed by post-hoc, immunohistochemial analyses of DREADD markers; hM4Di (GFP) and hM3Dq (mCherry) (**Figure 1D-E**).

### Chemogenetic Claustrum Inhibition Increases Amplitude of Spontaneous Activity

To causally test the effect of inhibiting claustrum activity on prefrontal cortex and other cortical networks, we transiently inactivated PFC-projecting claustrum cells by delivering CNO (5 mg/kg) to mice expressing hM4Di in claustrum cells while simultaneously recording mesoscale cortical activity Resting-state, spontaneous cortical activity was quantified activity by computing the root-mean-square (RMS) of regional, time-varying, spontaneous fluctuations in cortical fluorescence. Activity measurements from cortical ROIs were compared between three time points (Pre CNO, Post CNO, Post CNO (1.5h)). A mask was applied to the montage of spontaneous activity in **Figure 1F**.

ROIs were selected from experimentally derived cortical sensory mapping and extrapolations from the Allen Mouse Brain Atlas (Allen Reference Atlas – Mouse Brain; Oh et al., 2014) (**Figure 1G**). (**Figure 1G**). Following claustrum inhibition, we observed a prominent increase in spontaneous cortical activity in the medial anterior area near the retro-Cre injection location (PFC) but also within adjacent posterior-medial regions incorporating the cingulate cortex (CG) (**Figure 1I/J**). A two-way ANOVA was performed to analyze the ROIs and time points of the recordings relative to CNO injections on spontaneous activity measurements (RMS) in animals injected with hM4Di. Simple main effects analysis showed that ROIs and recording time points had a statistically significant effect on spontaneous activity **(Table 2)**. A statistically significant interaction between the ROIs and the time points of the recordings was also detected **(Table 2)**. Overall, inhibition of the claustrum using hM4Di resulted in widespread increased activity in medial frontal cortical regions encompassing the PFC, CG, and M2. Interestingly, there was also an increase in the PTA, V1 and RS (**Figure 1K**, **Table 2, Supplemental Table 1),** although increased activity in the RS was also found in the control condition (**Supplemental Figure 2**) and in response to the injection of saline (not shown), suggesting nonspecific increases across conditions. These increases in activity were significant in the Post CNO recording and most regions returned to near baseline levels at the Post CNO (1.5 hours) recording.

**Table 1.** Region-of-interest (ROI) Abbreviations and Locations.

|  |  |  |  |
| --- | --- | --- | --- |
| PFC | Prefrontal cortex (1) | UN | Unassigned (9) |
| CG | Cingulate (2) | S2 | Secondary somatosensory (10) |
| M2 | Secondary motor (3) | PTA | Parietal association cortex (11) |
| M1 | Primary motor (4) | RS | Retrosplenial cortex (12) |
| FL | Forelimb (5) | V1 | Primary visual cortex (13) |
| HL | Hindlimb (6) | AU | Auditory cortex (14) |
| BC | Barrel cortex (7) | TEA | Temporal association cortex (15) |
| TR | Trunk (8) |  |  |

**Table 2.**
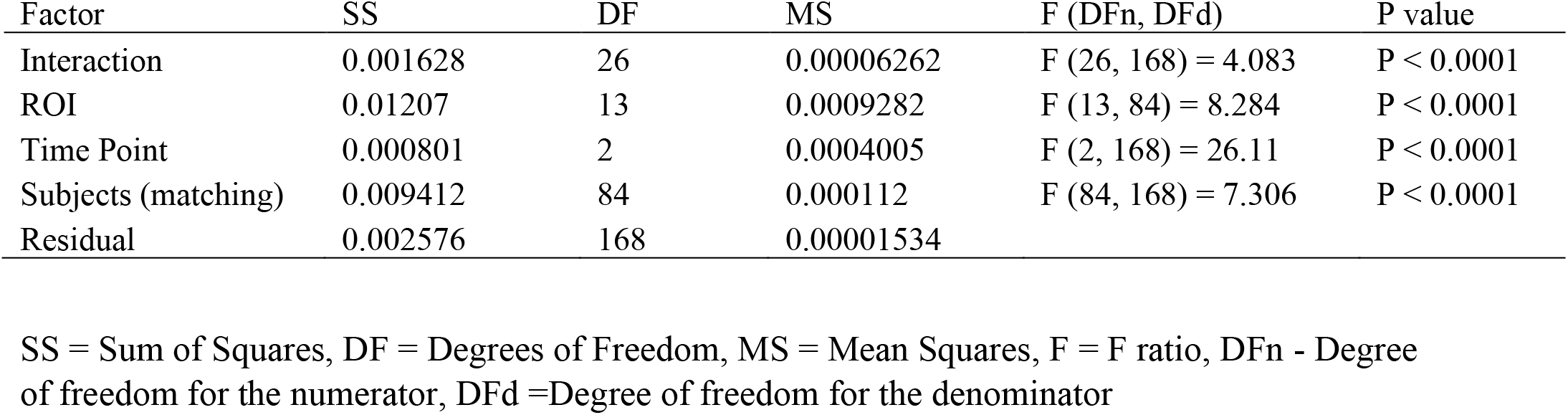
Spontaneous activity measurements from hM4Di mice, root-mean-square (RMS) ANOVA table.

### Chemogenetic Claustrum Excitation Decreases Amplitude of Spontaneous Activity

In a separate cohort of mice, we examined cortical activity following the excitation of claustrum neurons using hM3Dq (**Figure 2A**). In contrast to claustrum inhibition, activation of claustrum resulted in decreased activity in the PFC and a slight decrease in the CG (**Figure 2B-F**). Two-way ANOVA revealed main effects of the ROI locations and recording time points had a statistically significant effect on spontaneous activity **(Table 3)**. A statistically significant interaction between the ROIs and time points of the recordings was also detected **(Table 3)**. In contrast with hM4Di the effects on activity from hM3Dq was still present in the Post CNO (1.5h) recording (**Figure 2F**, **Table 3, Supplemental Table 2**).

**Figure 2.**
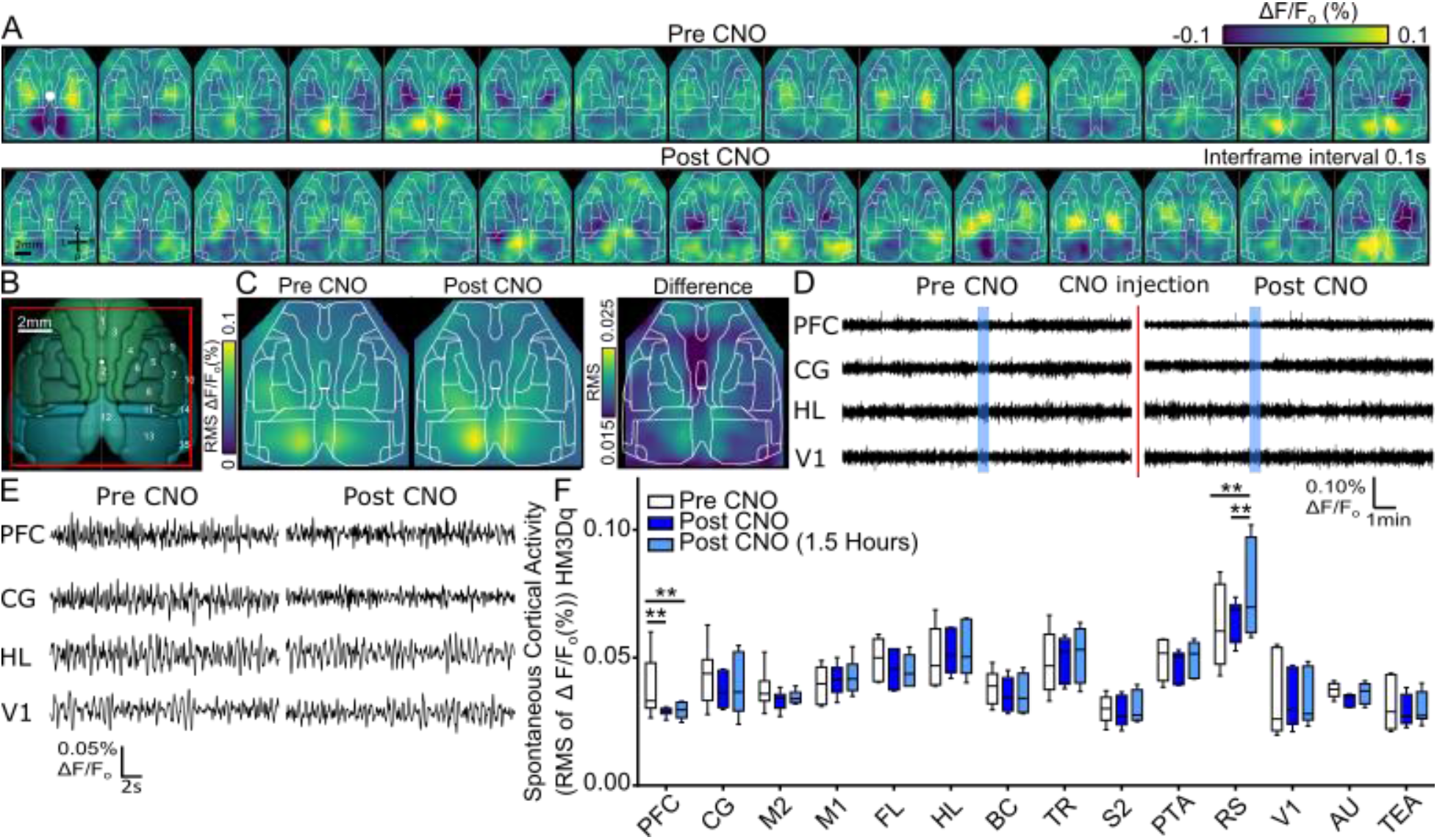
Chemogenetic excitation of claustrum activity by hM3Dq activation causes decreased spontaneous cortical activity. **A)** Montage of exemplar spontaneous cortical activity before (Pre) and after (Post) CNO injection (5 mg/kg i.p.). Inter-frame interval of 0.1 s, white dot denotes the bregma. **B)** Schematic of cortical regions of interest based on Allen Brain Institute 3D Brain Explorer, red box approximates imaging field of view. **C)** Mean cortical activity map of calculated root-mean-square (RMS) values across animals for each pixel on in imaging field of view before (left) and following (right) CNO treatment (5 mg/kg i.p.), n=7. Right panel shows the change in spontaneous activity (RMS) between the 2 conditions, where higher values indicated greater increases in RMS. **D)** Time course of a 15 minute recording from PFC, CG, HL, and V1 showing stable spontaneous activity over the duration of recording. Shaded blue areas indicate regions expand in Figure 2E. **E)** Example 30 s time-courses of spontaneous activity from PFC, CG, HL, and V1 before (Pre) and following (Post) CNO injection. **F)** Summary of changes in spontaneous cortical activity (n=6). Boxplot of RMS values calculated from the spontaneous cortical activity time course of the calcium signal for 14 ROIs spanning the entire cortical hemisphere. For each ROI, mean activity values were calculated for recordings prior to (white), immediately following (red), and 1.5 hours following (light red) CNO treatment (5 mg/kg i.p.). Statistical significance was calculated using a two-way ANOVA with a Tukey Test to correct for multiple comparisons. * P ≤ 0.05, ** P ≤ 0.01. Two-Way ANOVA with multiple groups comparison.

**Table 3.** hM3Dq root-mean-square (RMS ANOVA table.

| Factor | SS | DF | MS | F (DFn, DFd) | P value |
| --- | --- | --- | --- | --- | --- |
| Interaction | 0.001175 | 26 | 0.00004521 | F (26, 140) = 2.300 | P = 0.0011 |
| ROI | 0.02635 | 13 | 0.002027 | F (13, 70) = 10.10 | P < 0.0001 |
| Time Point | 0.0002071 | 2 | 0.0001035 | F (2, 140) = 5.268 | P = 0.0062 |
| Subjects (matching) | 0.01404 | 70 | 0.0002006 | F (70, 140) = 10.21 | P < 0.0001 |
| Residual | 0.002751 | 140 | 0.00001965 |  |  |

SS = Sum of Squares, DF = Degrees of Freedom, MS = Mean Squares, F = F ratio, DFn = Degree of freedom for the numerator, DFd = Degrees of freedom for the denominator

**Table 4.**
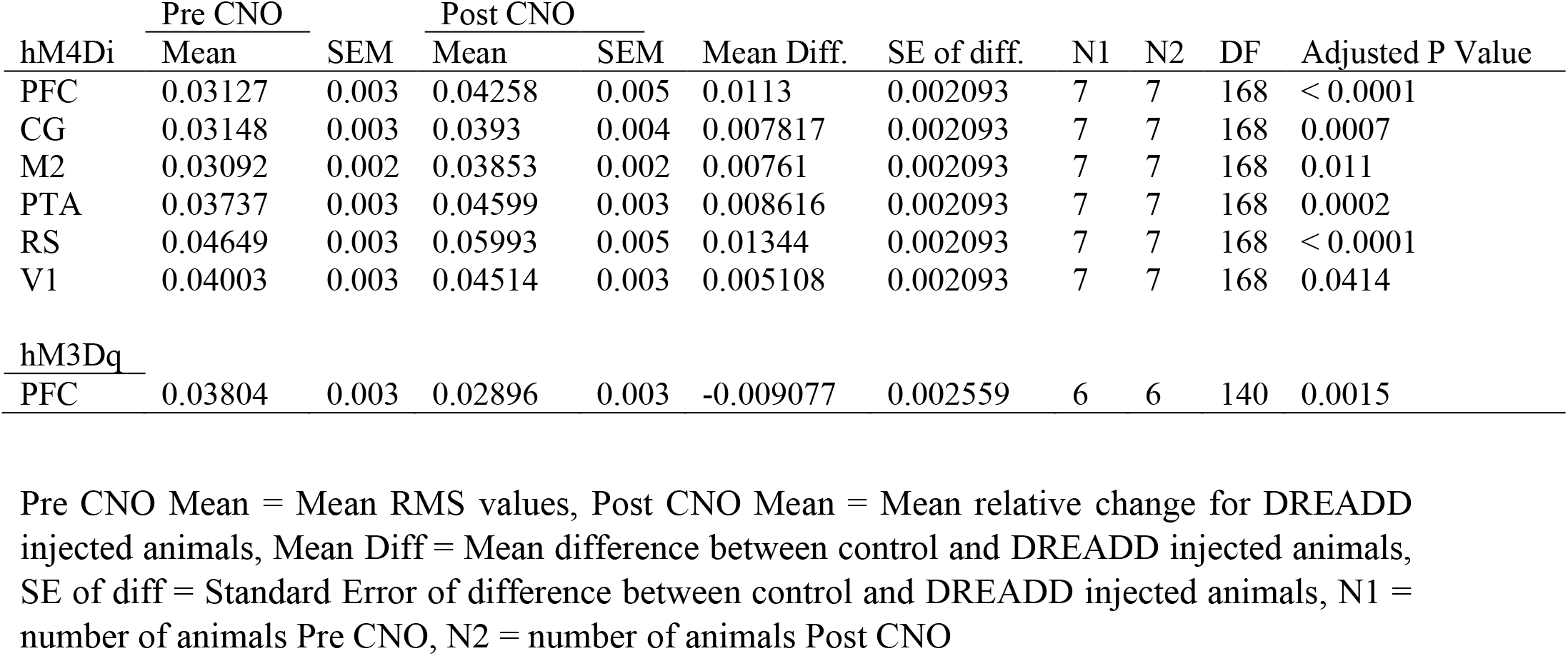
Summary of regional cortical activity changes between Pre and Post CNO recordings.

| hM4Di | Pre CNO |  | Post CNO |  | Mean Diff. | SE of diff. | N1 | N2 | DF | Adjusted P Value |
| --- | --- | --- | --- | --- | --- | --- | --- | --- | --- | --- |
|  | Mean | SEM | Mean | SEM |  |  |  |  |  |  |
| PFC | 0.03127 | 0.003 | 0.04258 | 0.005 | 0.0113 | 0.002093 | 7 | 7 | 168 | < 0.0001 |
| CG | 0.03148 | 0.003 | 0.0393 | 0.004 | 0.007817 | 0.002093 | 7 | 7 | 168 | 0.0007 |
| M2 | 0.03092 | 0.002 | 0.03853 | 0.002 | 0.00761 | 0.002093 | 7 | 7 | 168 | 0.011 |
| PTA | 0.03737 | 0.003 | 0.04599 | 0.003 | 0.008616 | 0.002093 | 7 | 7 | 168 | 0.0002 |
| RS | 0.04649 | 0.003 | 0.05993 | 0.005 | 0.01344 | 0.002093 | 7 | 7 | 168 | < 0.0001 |
| V1 | 0.04003 | 0.003 | 0.04514 | 0.003 | 0.005108 | 0.002093 | 7 | 7 | 168 | 0.0414 |
| hM3Dq |  |  |  |  |  |  |  |  |  |  |
| PFC | 0.03804 | 0.003 | 0.02896 | 0.003 | -0.009077 | 0.002559 | 6 | 6 | 140 | 0.0015 |

Pre CNO Mean = Mean RMS values, Post CNO Mean = Mean relative change for DREADD injected animals, Mean Diff = Mean difference between control and DREADD injected animals, SE of diff = Standard Error of difference between control and DREADD injected animals, N1 = number of animals Pre CNO, N2 = number of animals Post CNO

To ensure that the changes in spontaneous activity were specific to the manipulation of claustrum activity, we compared the relative change of RMS activity in hM4Di and hM3Dq to control animals (**Figure 3A**). A two-way ANOVA was performed to analyze the ROIs and time points of the recordings relative to CNO injections on spontaneous activity measurements (RMS) when comparing DREADD injected animals to the control animals. The two-way ANOVA revealed a statistically significant interaction between the effects of ROIs and time points of the recordings **(Table 5)**. Simple main effects analysis showed that ROIs and recordings time points all had a statistically significant effect on spontaneous activity **(Table 5)**.

**Figure 3.**
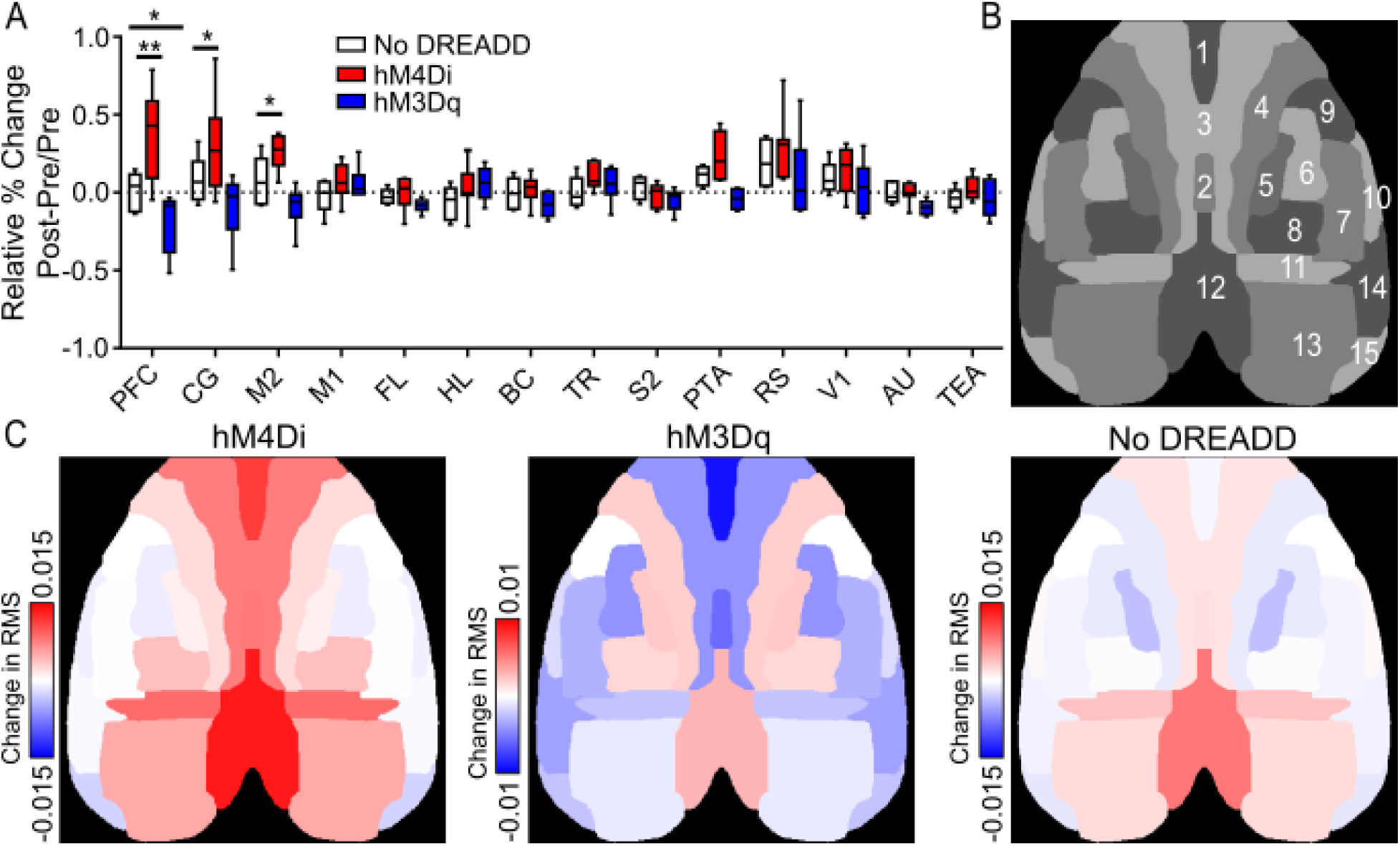
Summary of regional cortical activity changes resulting from chemogenetic manipulation of the claustrum. **A)** Summary of relative change in spontaneous cortical activity depicted as percent change of RMS for select ROIs comparing the Pre and Post CNO application time points between control animals (white, n=5), hM4Di (red, n=7), and hM3Dq (blue, n=6) animals. Statistics were calculated using a two-way ANOVA with a Dunnett’s Test to correct for multiple comparisons. * P ≤ 0.05, ** P ≤ 0.01. **B)** ROI labels for parcellated map. See Table 1 for ROIs represented by numbers. **C)** Mean change in RMS (ΔF/F˳) for animals injected with hM4Di (left. n=7), and hM3Dq (middle, n=6) and control animals (right, n=5).

**Table 5:** Relative change in RMS between no DREADD and hM4Di/hM3Dq ANOVA table.

| ANOVA table: |  |  |  |  |  |
| --- | --- | --- | --- | --- | --- |
| Relative Change | SS | DF | MS | F (DFn, DFd) | P value |
| Interaction | 13456 | 26 | 517.5 | F (26, 210) = 2.547 | P = 0.0001 |
| ROI | 10023 | 13 | 771 | F (13, 210) = 3.795 | P < 0.0001 |
| Condition | 11902 | 2 | 5951 | F (2, 210) = 29.29 | P < 0.0001 |
| Residual | 42667 | 210 | 203.2 |  |  |

SS = Sum of Squares, DF = Degrees of Freedom, MS = Mean Squares, F = F ratio, DFn = Degree of freedom for the numerator, DFd = Degrees of freedom for the denominator

**Table 6.**
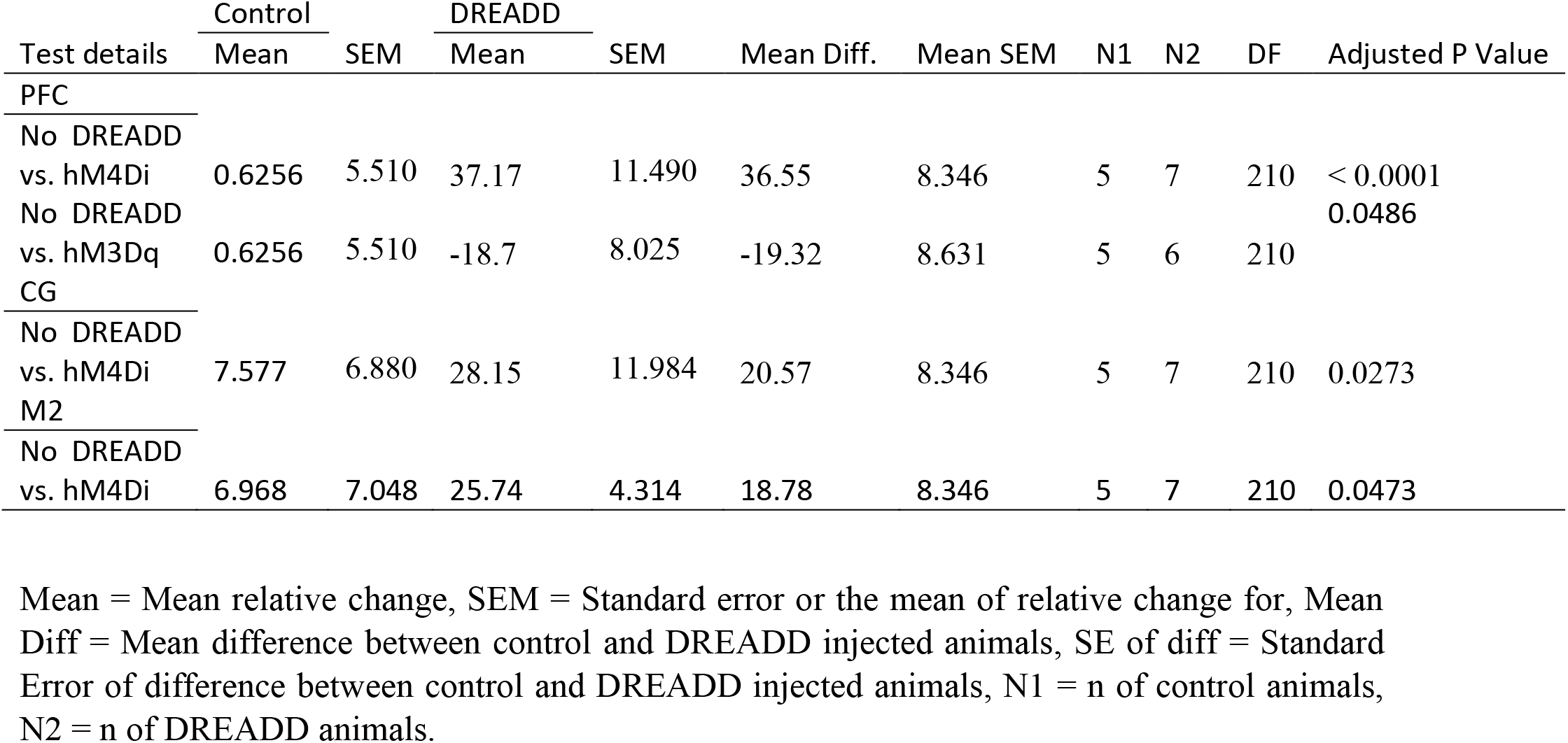
RMS relative change comparison between control and hM4Di/hM3Dq groups.

With hM4Di, the PFC, CG, and M2 regions showed significant increases in activity compared to the control animals, and HM3Dq showed a significant decrease in activity in the PFC (**Figure 3A**, **Table 5, Supplemental Table 5**). Changes in activity in off-target ROIs (PTA, RSC, V1) were not significant when compared to control animals. Therefore, chemogenetic claustrum manipulation results in specific changes in spontaneous activity in medial frontal cortical regions.

### Sensory Evoked Cortical Response

Next, we probed the impact of our manipulation of claustrum activity on cortical sensory-evoked processing following stimulation of different sensory modalities. Visual stimulation was delivered via full-field LED light flash delivered to the right eye (**Figure 4A**). Chemogenetic claustrum inhibition increased the peak amplitude of the sensory-evoked response but did not affect the kinetics of the response to sensory stimulation in the primary visual cortex (**Figure 4B-E**, Left). Hindlimb somatosensory stimulation was delivered via a 0.4s 10Hz stimulation from a piezoelectric bender delivered to the right hindlimb (**Figure 4C**). The kinetics or amplitude of the response to this stimulation were not altered following claustrum inhibition. Similarly, there was no change in the sensory-evoked visual or hindlimb stimulation in mice after claustrum activation or in CNO control mice. Therefore, across the sensory modalities examined, visual stimulation was the only sensory paradigm that showed a change with chemogenetic intervention.

**Figure 4.**
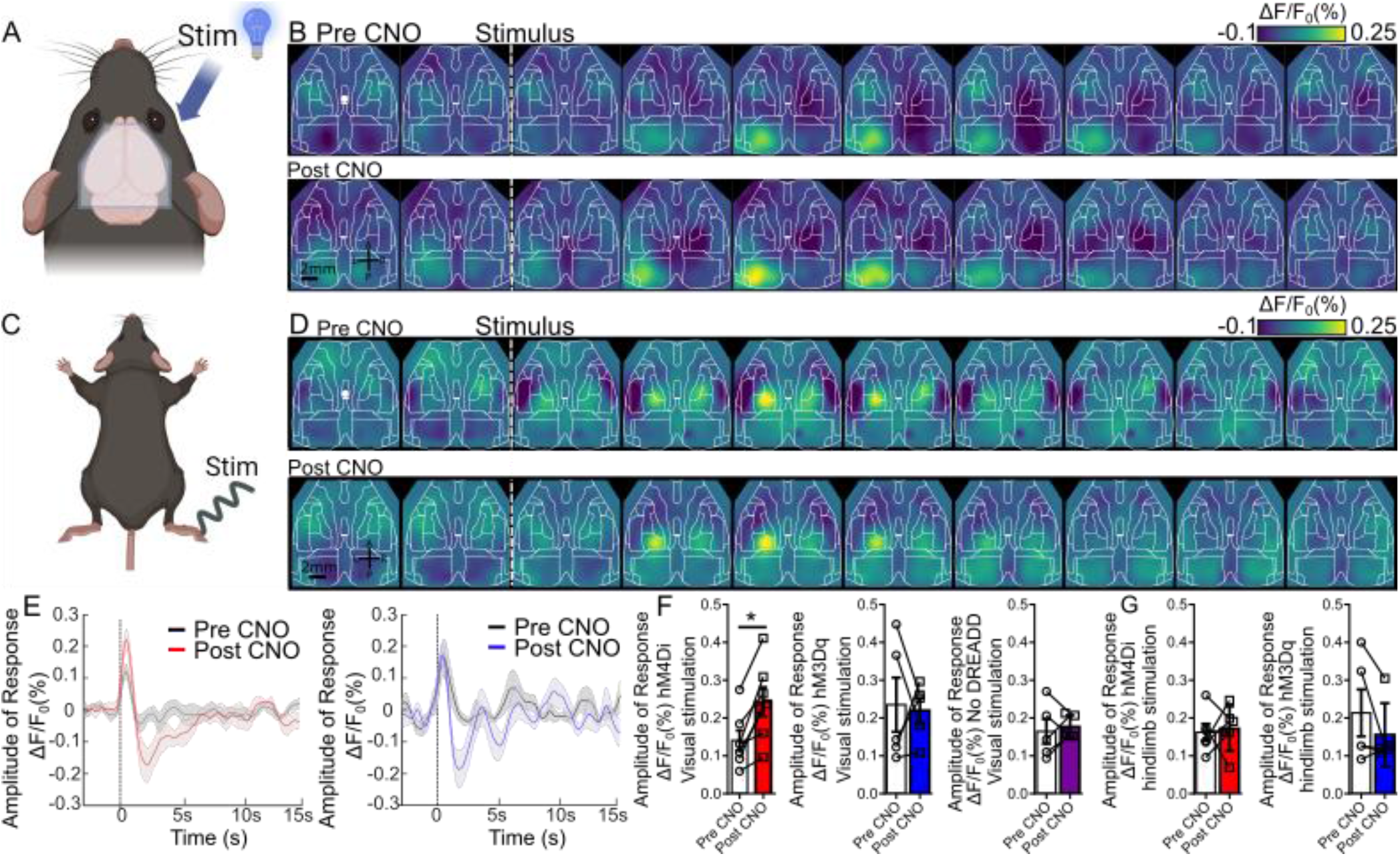
Claustrum inhibition increases amplitude of visual-evoked cortical response. **A)** Schematic of visual sensory stimulation; Visual stimulation delivered as a 3.5 ms blue light flash to the right eye. Created with BioRender.com **B)** Montage of averaged visual stimulation-evoked cortical activity in mice injected with hM4Di before (Pre) and after (Post) CNO injection. Inter-frame interval of 0.2 s. White circle denotes location of the bregma. **C)** Schematic of hindlimb sensory stimulation delivered via piezoelectric actuator to the right hindlimb. Created with BioRender.com **D)** As in B, montage of hindlimb sensory-evoked cortical activity in mice injected with HM4Di before (Pre) and after (Post) CNO injection **E)** Mean time courses for visual-evoked cortical activity for hM4Di (left, n=7) and hM3Dq (right, n=5) injected animals. Pre CNO condition is shown in Black and Post CNO in red for hM4Di and blue for hM3Dq. Shaded areas are standard error of the mean. **F)** Peak value of sensory responses for visual sensory evoked cortical responses Pre and Post CNO treatment with mice injected with hM4Di (left, n=7), hM3Dq (centre, n=5), and control animals (right, n=7) Error bars indicate SEM. * P ≤ 0.05 **G)** Peak value of sensory-evoked cortical responses for hindlimb stimulation Pre and Post CNO treatment with mice injected with hM4Di (Left, n=6), hM3Dq (Right, n=5). Error bars indicate SEM.

### Increased Local Connectivity with Claustrum Inhibition

We assessed changes in local connectivity resulting from claustrum manipulation using seed pixel analysis. We generated seed pixel correlation maps by using the same coordinate ROIs determined previously (**Figure 5A**). Claustrum inhibition using hM4Di resulted in an increased area of local connectivity in the same regions that showed increased activity, the PFC and CG, indicating that the observed increased activity was synchronized over broad adjacent topographical regions. Conversely, a trend toward decreased recruitment of local areal connectivity was observed when claustrum activity was excited with hM3Dq (**Figure 5C**).

**Figure 5.**
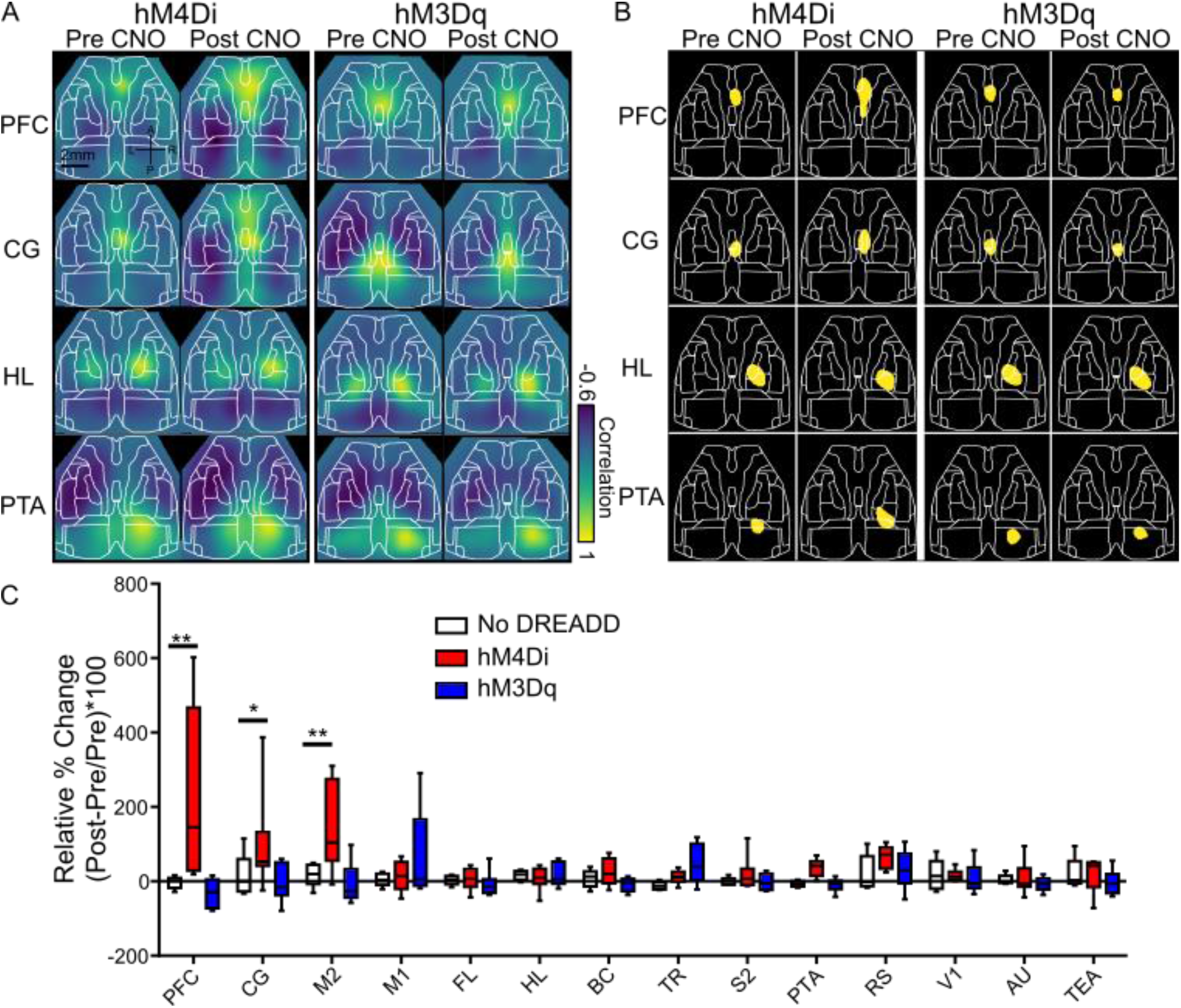
Chemogenetic claustrum inhibition via hM4Di increases local functional connectivity in PFC and medial cortices. **A)** Seed pixel maps of PFC, CG, HL, and PTA from the right hemisphere before and following CNO administration in hM4Di (Left) and HM3Dq mice (Right). **B)** Area changes for each ROI in panel A based on a threshold for consistent baseline area. **C)** Relative change for each ROI compared the area in the Pre CNO recording for hM4Di (red, n=7), hM3Dq (blue, n=6) and control mice (white, n=5). All measurements are from right hemisphere ROIs. Whiskers indicate max and min ranges, box 25^th^-75^th^ percentiles, and the line in the box indicates median value. Black line indicates baseline area. Statistics were calculated using a two-way ANOVA with a Dunnett’s Test to correct for multiple comparisons. * P ≤ 0.05, ** P ≤ 0.01. PFC (F (13, 210) = 1.691, p=<0.0001), CG (F (13, 210) = 1.691, p=0.0339), M2 (F (13, 210) = 1.691, p=0. 0033).

**Figure 6.**
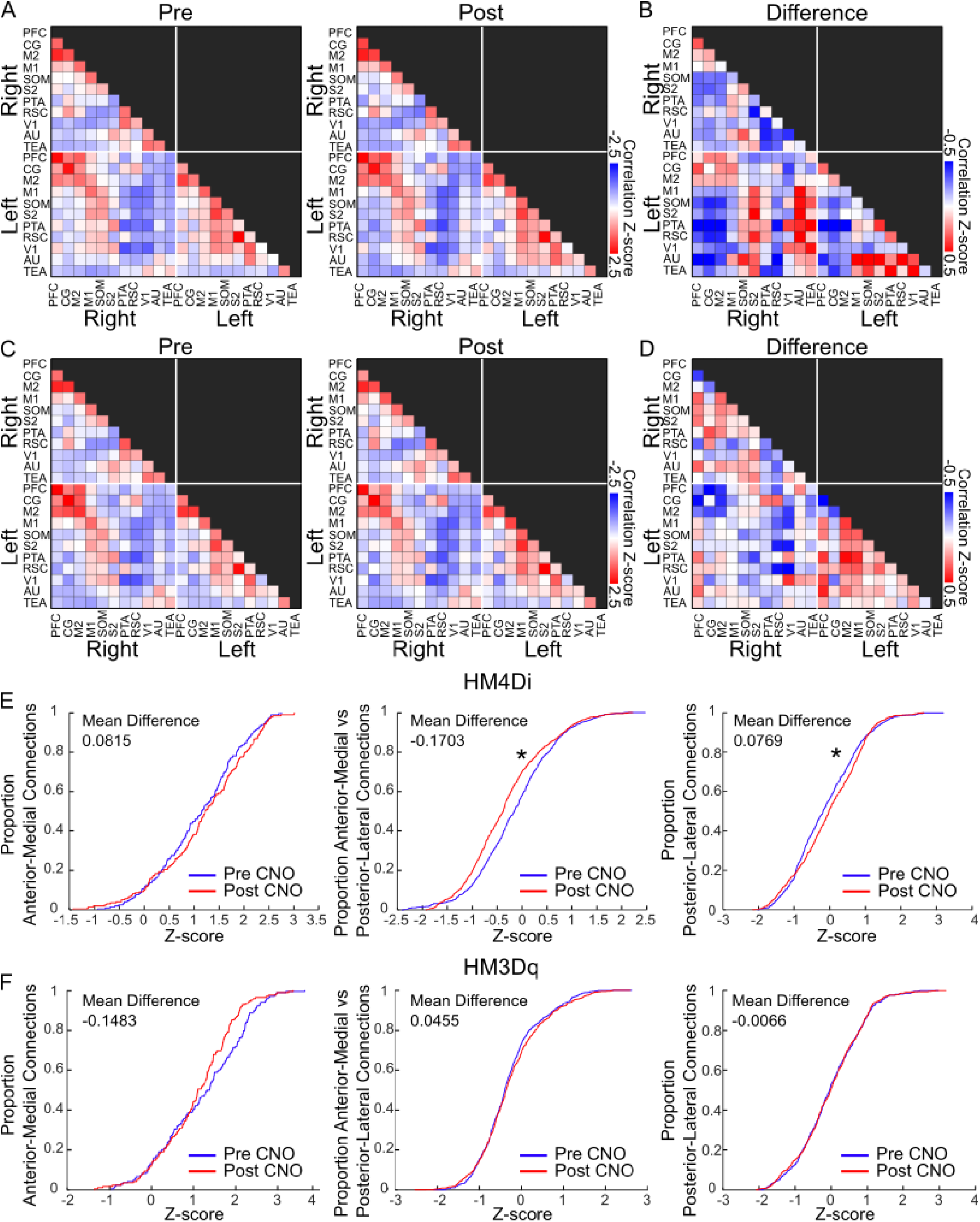
Claustrum inhibition induces decoupling of functional connectivity between long-range cortical regions. **A)** Functional connectivity matrix of mean z-score correlation values derived from spontaneous activity from hM4Di (n=7) Pre CNO (Left), and Post CNO (Right). **B)** Difference matrix calculated between mean hM4Di Post CNO and Pre CNO z-score correlation matrices. **C)** Mean z-score correlation matrix from hM3Dq (n=6) Pre CNO (Left), and Post CNO (Right). **D)** Difference matrix calculated between mean hM3Dq Post CNO and Pre CNO z-score correlation matrices. **E)** Cumulative distribution function for connections (n=7) within the anterior-medial ROIs (Left), between anterior-medial and posterior-lateral ROIs, and within posterior-lateral ROIs (Right) in hM4Di injected mice. Statistics were calculated using a Wilcoxon rank sum test. **F)** Cumulative distribution function for connections (n=6) within the anterior-medial ROIs (Left), between anterior-medial and posterior-lateral ROIs, and within posterior-lateral ROIs (Right) in hM3Dq injected mice.

### Inter-regional Functional Connectivity Changes

Functional connectivity changes within and between cortical regions were examined by computing region-of-interest-based correlation matrices for each mouse from the Pre CNO and Post CNO spontaneous recordings. Baseline, Pre CNO correlation matrices for hM4Di and hM3Dq injected animals showed consistent patterns of functional connectivity (**Figure 6A**, **Supplemental Figure 4)**. To visualize the changes between Pre and Post CNO we calculated a difference matrix by subtracting the z-score values of the Pre CNO matrix from the Post CNO matrix (**Figure 6B**). The difference matrix from animals with claustrum inhibition reveal broad cortex-wide alterations in regional functional connectivity. In particular, two broad themes of increased connectivity among anterior-medial regions, and decreased connectivity between anterior-medial and posterior-lateral regions. When we examined the z-score matrices Pre and Post CNO and animals injected with hM3Dq (**Figure 6C**) and calculated the difference matrix (**Figure 6D**) we observed the corollary trend wherein anterior-medial regions exhibited decreased connectivity, and increased connectivity between anterior-medial and posterior-lateral regions.

To quantify the difference between Pre and Post CNO matrices we grouped ROIs into 2 categories, anterior-medial regions which included PFC, CG, M1, and M2, and posterior-lateral regions which included the other ROIs. From these groupings, we calculated connectivity cumulative distribution functions (CDFs) within and between the anterior-medial and posterior-lateral regional categories. There was a further reduction in the correlation between the anterior-medial and posterior-lateral groups (**Table 7**), but no change in inter-regional or intra-region coupling following claustrum excitation or in control mice (**Figure 6E** and **Supplementary Figure 5**).

**Table 7.** Global functional connectivity changes.

| hM4Di | Pre CNO |  | Post CNO |  | P value |
| --- | --- | --- | --- | --- | --- |
|  | Mean | SEM | Mean | SEM |  |
| Anterior-medial vs anterior-medial | 1.11 | 0.059 | 1.18 | 0.065 | p =0.2314 |
| Anterior-medial vs posterior-lateral | -0.11 | 0.037 | -0.03 | 0.037 | p =0.0482 |
| Posterior-lateral vs posterior-lateral | -0.16 | 0.028 | -0.33 | 0.028 | p =0.009 |
| **hM3Dq** |  |  |  |  |  |
| Anterior-medial vs anterior-medial | 1.23 | 0.077 | 1.08 | 0.069 | p =0.1121 |
| Anterior-medial vs posterior-lateral | -0.31 | 0.028 | -0.26 | 0.30 | p =0.6241 |
| Posterior-lateral vs posterior-lateral | -0.018 | 0.038 | -0.0485 | .0039 | p =0.2559 |

Mean = Mean CDF, SEM = Standard error or the mean of the CDF

## Discussion

We assessed the functional role of claustrocortical projections to cortex using projection-specific, chemogenetic modulation of claustrum activity along with the simultaneous assessment of cortex-wide dynamics via mesoscale fluorescence imaging. Inhibition of claustrum resulted in increased activity in PFC, consistent with previous reports indicating claustrum has a feed-forward inhibitory function on cortical activity (Atlan et al., 2018; Jackson et al., 2018; Chia et al., 2020; Norimoto et al., 2020; McBride et al., 2023). However, we also observed profound increases in spontaneous activity in adjacent lateral regions, including M2, and medio-posterior regions, including the cingulate cortex, indicating a more expansive functional impact for PFC projecting claustrum neurons. That we observed parallel but opposing region-specific changes when we manipulated claustrum activity via hM3Dq activation, leads credence to these observations and suggests a resting-state claustrocortical tone of activity that facilitates bidirectional control of widespread cortical activity.

This is the first demonstration implicating bidirectional claustrocortical control of expansive resting-state cortical activity. Importantly, the pattern of altered functional cortical activity is supported by the anatomical topographic organization of claustrum projections projecting to the cortex (Marriott et al., 2020; Ham and Augustine, 2022; Wang et al., 2023).

The PFC is involved in many high-order processes, such as attention, decision-making, and memory, and is a critical component of the default mode network whose activity predominates during resting-states (Smith et al., 2019). Therefore, the ability of the claustrum to bidirectionally modulate activity in the PFC has significant implications for the potential function of the claustrum. The observation that M2 and CG also exhibited altered activity may indicate a parallel control pathway from claustrum to coordinate activity in these adjacent areas. Claustral neurons projecting to the PFC also exerting inhibition to surrounding areas that also have a role in higher-order processes highlights the strength of the feedforward inhibitory control the claustrum has over frontal cortical regions, and the effects on M2 suggest that the claustrum may have a role in planning motor behaviours.

The claustrum has dense reciprocal connections to sensory cortices in rodents (Torgerson et al., 2011; Wang et al., 2017; Wang et al., 2023) and has been speculated to be involved in salience detection (Jackson, 2018; Jackson et al., 2020; Madden et al., 2022), multisensory integration, and perceptual binding (Smith et al., 2019; Remedios et al., 2010). We found that claustrum inhibition increased the visual-evoked cortical responses. Indeed, discrete visual subdivisions of claustrum connected to primary visual areas and retaining retinotopic features have been described previously (Olson & Graybiel, 1980) and recent systematic viral tracing studies have revealed dense, bidirectional connections between primary visual cortex and ipsilateral claustrum (Wang et al., 2017). We previously described a topographic organization of claustrum projections, whereby anterior claustrum neurons project to anterior cortical regions, and posterior claustrum neurons project to poster cortical regions (Marriott et al., 2020) and as such it is unclear if the claustrocortical projections targeted in this study are directly modifying activity within primary visual cortex. Zhang et al. (2014) showed that the CG has long-range cortical projections to the visual cortex, and that optogenetic activation of CG neurons increased the response in visual cortical neurons during both anesthetized and awake states (Zhang et al., 2014). In a follow up paper, Zhang et al. (2016) found that the anterior cingulate area has reciprocal connections to the visual cortex, and a more recent study showed that optogenetic stimulation of anterior cingulate neurons activated VIP interneurons in the visual cortex, enhancing performance visual tasks, and this effect was prevented by chemogenetically inhibiting VIP interneurons in the visual cortex (Bastos et al., 2023). There is also research showing that the anterior cingulate cortex has a claustro-cortical connection whereby anterior cingulate signals are propagated to the visual cortex through the claustrum (White and Mathur, 2017). Thus, the enhancement of visual-evoked activity may be a downstream consequence of claustrum-mediated disinhibition of these modulatory cingulate areas.

Inhibition of the claustrum resulted in profound local and remote changes in cortical functional connectivity. When the claustrum was inhibited, seed pixel correlation analysis revealed a profound expansion of areal PFC, indicating that the increased spontaneous activity observed in Figure 1 was synchronized and spatially expansive. Remarkably, expanded local connectivity was similarly observed in adjacent regions including M2 and CG. A similar expansion of local connectivity was not observed in control mice nor in conditions where claustrum activity was increased via hM3Dq. This may result from the differing mechanism of action of hM3Dq activity in neurons wherein excitability alterations are mediated by Gq versus Gi-mediated G-protein signalling (Armbruster et al., 2007; Alexander et al., 2009; Roth, 2016; Atasoy & Sternson, 2018). Our region-based correlation analysis revealed even broader alterations in cortex-wide functional connectivity resulting from the manipulation of claustrum activity. Consistent with seed-pixel analyses, we observed increased inter-regional connectivity between areas of enhanced activity between PFC, M2, and CG. When considered as a regional module, we observed a decoupling of connectivity with these anterior-medial regions with posterior-lateral regions including SOM, S2, PTA, and RS, when the claustrum was inhibited. This suggests that the claustrum normally facilitates the interactions between these distant cortical networks, and a reduction in claustrum activity enables these spatially and functionally dissociable cortical regions to function more autonomously. This observed result is in accordance with the proposed role of the claustrum in cortical control (Mitra and Raichle, 2016; Wright et al., 2017; Madden et al., 2022).

Recently, several papers have measured changes in cortical activity following claustrum manipulation. One recent paper (Lou et al., 2022) used DREADDs to excite claustrum neurons and showed decreased low-frequency field potential activity in the cortex. However, their approach activated all claustrum neurons, without pathway specificity, so it is unclear which claustrum projections are responsible for the observed effects. Another recent paper (McBride et al., 2023) used optogenetics to modulate claustrum activity and showed a cortical layer and brain region-specific activation or inhibition of neural activity. However, no pathway-specific activation was used, and the electrophysiological approach employed limited recordings to one region at a time. In contrast, our study used pathways-specific manipulation and widefield imaging to attempt to isolate the effect of manipulating the claustrum projection to the prefrontal cortex. The large imaging window we employed allowed us to measure nearly the entire cortex to observe global cortical changes and changes in connectivity that would be more difficult to observe using electrophysiology. In the future, this method could be used to target other claustrum output streams (such as to the medial and lateral entorhinal cortex) to compare how these separate claustrum projections control cortical activity and inter-regional communication.

There have been several proposed models of claustrum function (Smith et al., 2019; Madden et al., 2022), but there are still many gaps in the understanding of how the claustrum interacts with the cortex at a global level. One aspect of the cognitive control model (Madden et al., 2022) proposes region-specific effects of claustral cortical projections that are consistent with our findings. These large-scale organizational control mechanisms may contribute to observed claustrum-mediated abnormalities associated with neuropsychiatric disorders relating altered claustrum activity (Galeno et al., 2009), reduced claustrum anterior cingulate connectivity (Jacobson-McEwen et al., 2014) in patients with schizophrenia, and reduced connectivity between the claustrum and the anterior cingulate in some cases of MDD (Bernstein et al., 2016). Given the elongated and topographical organization of both inputs and outputs of the claustrum, different claustrum subregions may serve different functions. Therefore, it is important to tease out the function of different claustrum subpopulations and to observe their effects on different cortical regions. With the difficulty of specifically manipulating claustrum activity by conventional methods, the use of pathway-specific techniques is vital for further understanding the function of the claustrum. Our results show pathway-specific claustrum excitation and inhibition result in opposing effects on local and long-range coordination of activity in cortical networks. The results of this study help further the understanding of how the claustrum influences cognition through distinct effects on cortical networks.

## Conflict of interest statement

The authors have nothing to declare.

## Acknowledgements

Natural Sciences and Engineering Research Council of Canada, Grant/Award Number: RGPIN202-05988 (AWC). Brain Canada Future Leaders Award (AWC). Scottish Rite Charitable Foundation Puzzles of the Mind Award (AWC). Canadian Institutes of Health Research, Grant/Award Number: 426485 (JJ). National Alliance for Research on Schizophrenia and Depression (JJ). Natural Sciences and Engineering Research Council of Canada, Grant/Award Number: RGPIN2018-05212 (JJ).

## Supplemental Figures

**Supplemental Figure 1:**
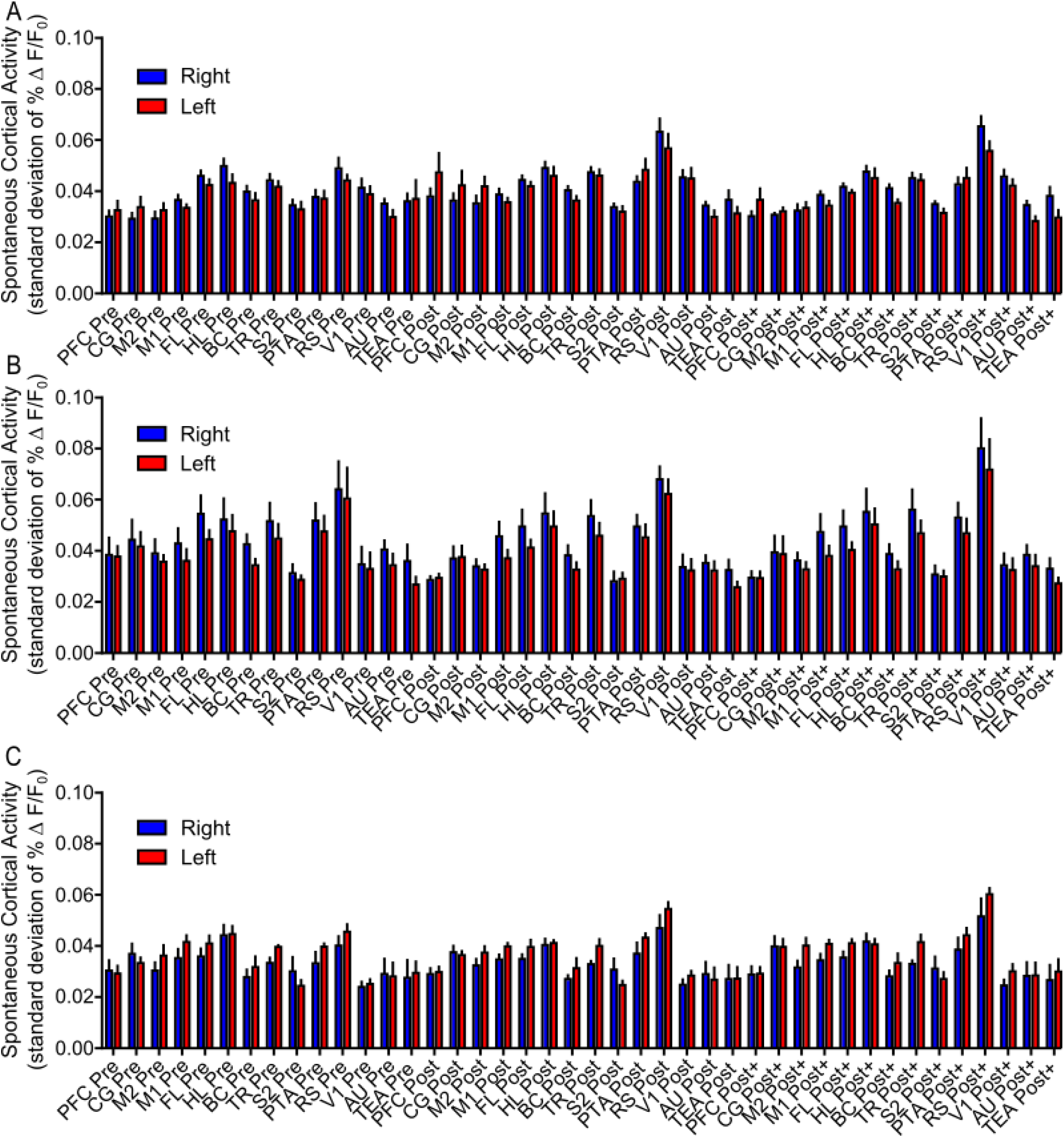
Pre CNO spontaneous activity RMS values from ROIs are similar in each cortical hemisphere. **A)** Bar chart comparing RMS values from Pre CNO recordings between left and right hemispheres in hM4Di injected animals (n=7). No significant differences were observed between any homotopic brain regions. Statistics were done using a two-way ANOVA with Bonferroni’s Test to correct for multiple comparisons. **B) As in A,** bar chart comparing RMS values from Pre CNO recordings between left and right hemispheres in hM3Dq injected animals (n=6). No significant differences were observed between any homotopic brain regions. Statistics were done using a two-way ANOVA with Bonferroni’s Test to correct for multiple comparisons. **C) As in A and B,** bar chart comparing RMS values from Pre CNO recordings between left and right hemispheres in control animals (n=5). No significant differences were observed between any homotopic brain regions. Statistics were done using a two-way ANOVA with Bonferroni’s Test to correct for multiple comparisons.

**Supplemental Figure 2:**
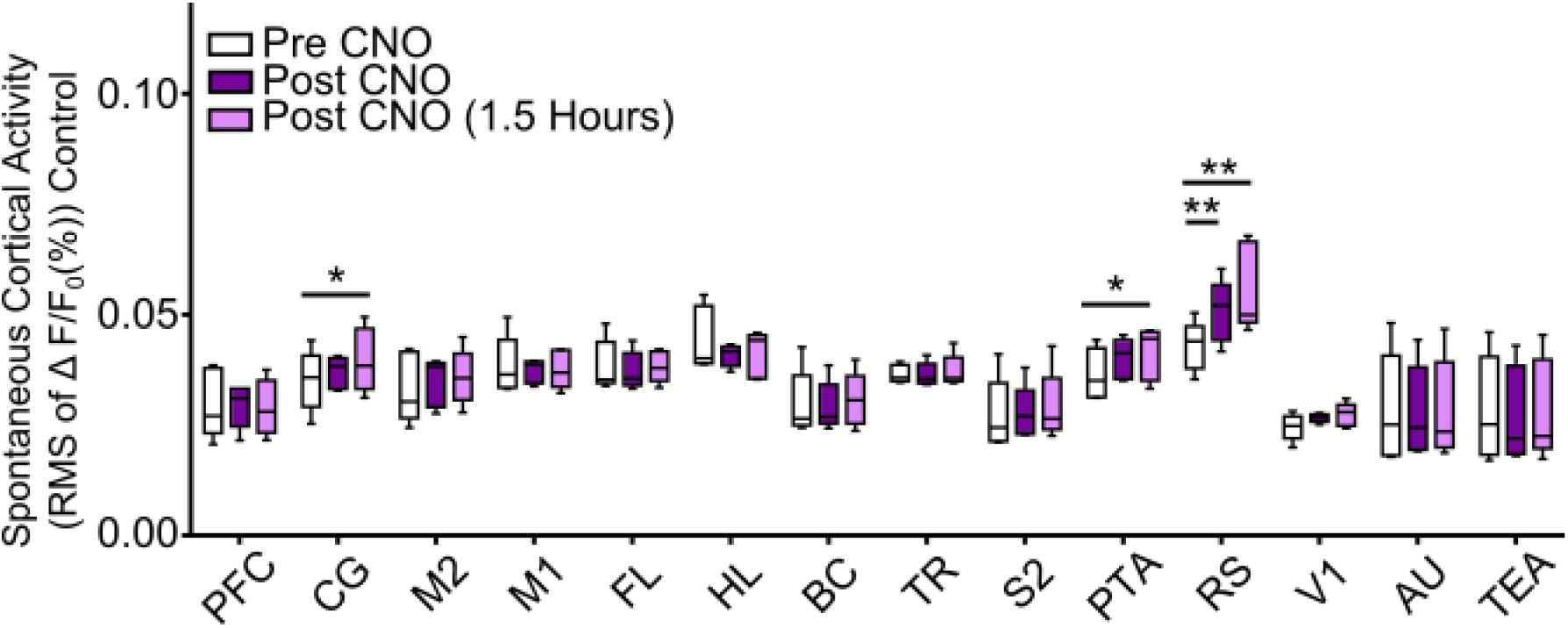
RMS values in control mice. Boxplot of RMS calculated from the spontaneous cortical activity time course of the calcium signal for 14 ROIs in control mice receiving CNO but without DREADDs. For each ROI, mean activity values were calculated for recordings prior to (black), immediately following (red), and 1.5 hours following (blue) CNO treatment (5mg/kg i.p.). Whiskers indicator max and min ranges, box 25^th^-75^th^ percentiles, and the line in the box indicates median value. Statistics were done using a two-way ANOVA with a Tukey Test to correct for multiple comparisons. Only RS showed increased activity between Pre CNO and Post CNO recordings (RS_Pre CNO_ 0.043±0.002, RS_Post CNO_ 0.051±0.003). * P ≤ 0.05, ** P ≤ 0.01. Two-Way ANOVA with multiple groups comparison (n=5).

**Supplemental Figure 3.**
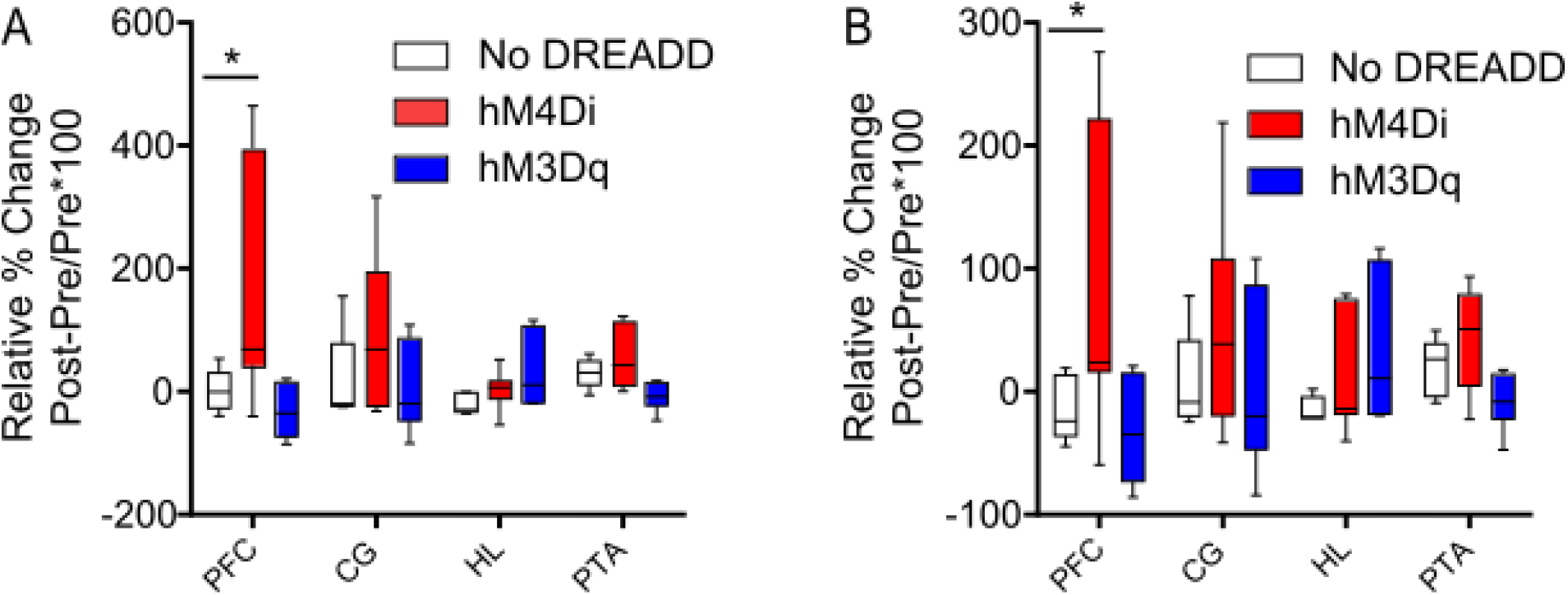
Seed pixel map areal changes at different thresholds. **A)** Relative change for each ROI compared the area in the Pre CNO recording for hM4Di (red), HM3Dq (blue) and control mice (white) at a threshold of 98^th^ percentile (99^th^ percentile for HL). All measurements are from right hemisphere ROIs. Whiskers indicate max and min ranges, box 25^th^-75^th^ percentiles, and the line in the box indicates median value. Black line indicates baseline area. Statistics were calculated using a two-way ANOVA with a Dunnett’s Test to correct for multiple comparisons. PFC Pre CNO 1.30±15.33, PFC Post CNO 150.13±73.98. * P ≤ 0.05. PFC (F (2, 60) = 6.550 p=0.0077). **B)** Relative change for each ROI compared the area in the Pre CNO recording for hM4Di (red), hM3Dq (blue) and control mice (white) at a threshold of 95^th^ percentile. All measurements are from right hemisphere ROIs. Whiskers indicate max and min ranges, box 25^th^-75^th^ percentiles, and the line in the box indicates median value. Black line indicates baseline area. Statistics were calculated using a two-way ANOVA with a Dunnett’s Test to correct for multiple comparisons. PFC_Pre CNO_ −13.60±11.80, PFC_Post CNO_ 79.65±46.02. * P ≤ 0.05. PFC (F (2, 60) = 5.141, p=0.0231).

**Supplemental Figure 4.**
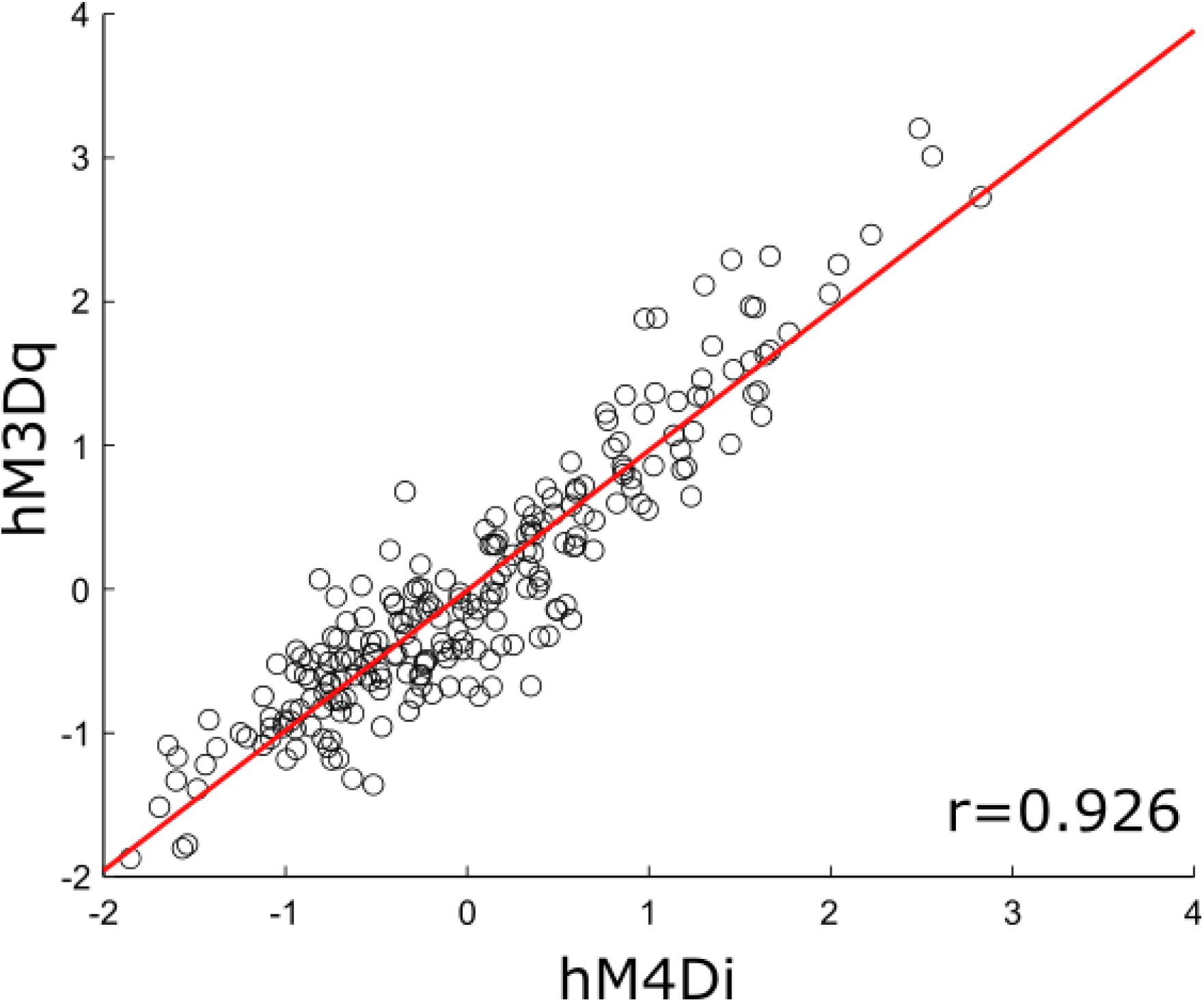
Pre CNO correlation matrix values similar between hM4Di and hM3Dq injected animals. Scatter plot of z-score correlation between hM4Di- and hM3Dq-injected animals Pre CNO correlation matrices with line of best (red line). Correlation was calculated using a linear correlation, r=0.926.

**Supplemental Figure 5.**
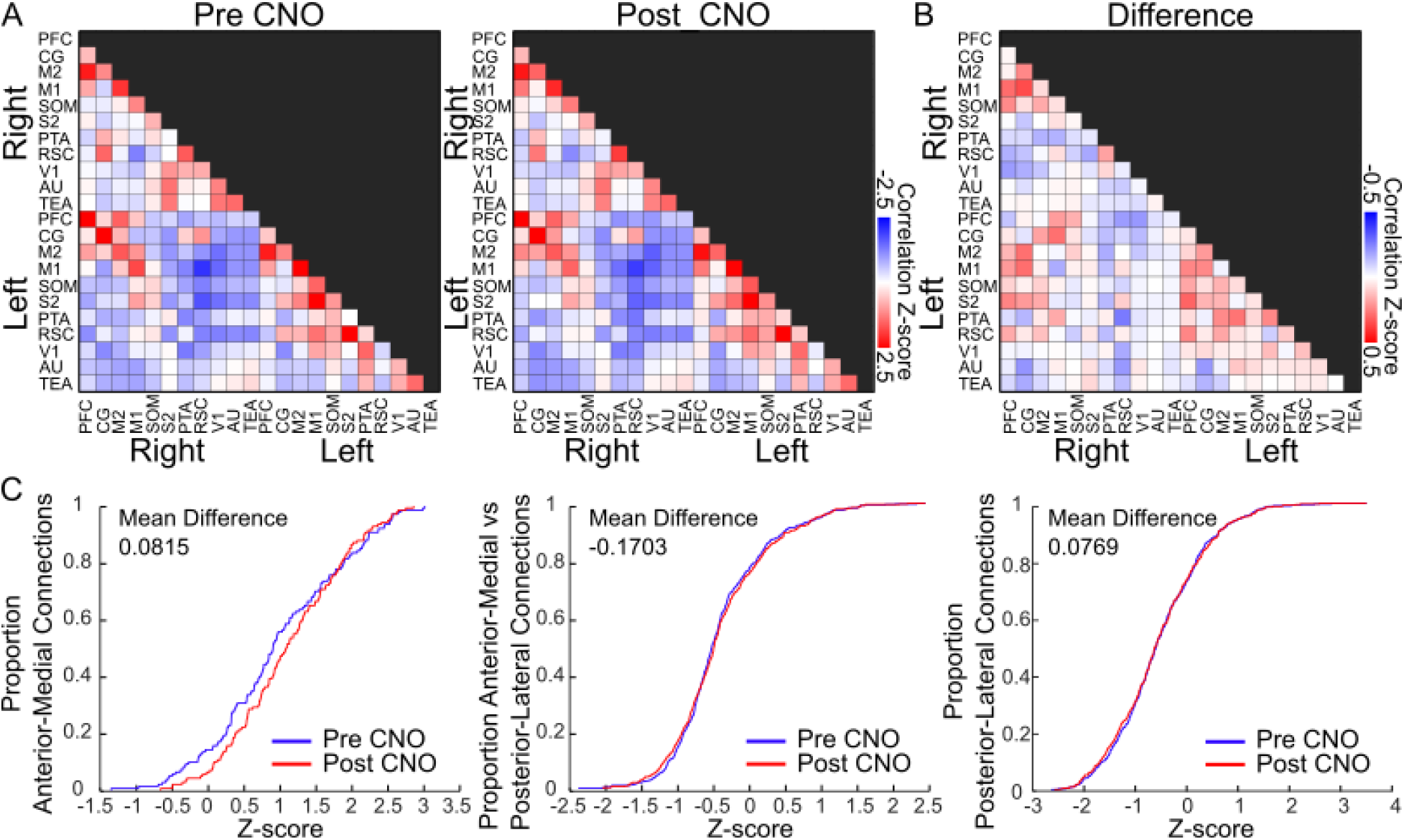
No effect on global cortical dynamics in control animals without DREADDs. **A)** Mean z-score correlation matrix from control animals, Pre CNO (Left), and Post CNO (Right). **B)** Difference between mean Post CNO and Pre CNO z-score correlation matrices in control animals. **C)** Cumulative distribution function for connections within the anterior-medial ROIs (Left), between anterior-medial and posterior-lateral ROIs, and within posterior-lateral ROIs (Right) in control mice. Statistics were calculated using a Wilcoxon rank sum test.

## Notes

### Competing Interest Statement

The authors have declared no competing interest.

